# Recapitulating the life cycle of the global pathogen *Entamoeba* in mice

**DOI:** 10.1101/2022.10.26.513970

**Authors:** Carolina Mendoza Cavazos, Marienela Y. Heredia, Leah A. Owens, Laura J. Knoll

## Abstract

There are several *Entamoeba* species that colonize humans, but only *Entamoeba histolytica* causes severe disease. *E. histolytica* is transmitted through the fecal-oral route to colonize the intestinal tract of 50 million people worldwide. The current mouse model to study *E. histolytica* intestinal infection directly delivers the parasite into the surgically exposed cecum, which circumvents the natural route of infection and does not produce infectious cysts. To develop a fecal-oral mouse model, we screened our vivarium for a natural murine *Entamoeba* colonizer via a pan-*Entamoeba* PCR targeting the 18S ribosomal gene. We determined that C57BL/6 mice were chronically colonized by *Entamoeba muris*. This amoeba is closely related to *E. histolytica*, as determined by 18S sequencing and cross-reactivity with an *E. histolytica*-specific antibody. In contrast, outbred Swiss Webster (SW) mice were not chronically colonized by *E. muris*. We orally challenged SW mice with 1×10^5^ *E. muris* cysts and discovered they were susceptible to infection, with peak cyst shedding occurring between 5-7 days post-infection. Most infected SW mice did not lose weight significantly but trended toward decreased weight gain throughout the experiment when compared to mock-infected controls. Infected mice treated with paromomycin, an antibiotic used against non-invasive intestinal disease, do not become colonized by *E. muris*. Within the intestinal tract, *E. muris* localizes exclusively to the cecum and colon. Purified *E. muris* cysts treated with bovine bile *in vitro* excyst into mobile, pre-trophozoite stages. Overall, this work describes a novel fecal-oral mouse model for the important global pathogen *E. histolytica*.

**Importance:** Infection with parasites from the *Entamoeba* genus are significantly underreported causes of diarrheal disease that disproportionally impact tropical regions. There are several species of *Entamoeba* that infect humans to cause a range of symptoms from asymptomatic colonization of the intestinal tract to invasive disease with dissemination. All *Entamoeba* species are spread via the fecal-oral route in contaminated food and water. Studying the life cycle of *Entamoeba*, from host colonization to infectious fecal cyst production, can provide targets for vaccine and drug development. Because there is not an oral challenge rodent model, we screened for a mouse *Entamoeba* species and identified *Entamoeba muris* as a natural colonizer. We determine the peak of infection after an oral challenge, the efficacy of paromomycin treatment, the intestinal tract localization, and the cues that trigger excystation. This oral infection mouse model will be valuable for the development of novel therapeutic options for *Entamoeba* infections.

## Introduction

Parasitic diseases are underappreciated causes of morbidity and mortality because disease outcomes are variable (1), cases are often underreported (2), and disease disproportionately impacts geographical locations experiencing poverty (1–3). Diarrheal diseases are a significant and underreported cause of child mortality in tropical regions (4, 5). Such infections are exacerbated by factors such as resource availability, lack of sanitation infrastructure, and malnutrition (6). For these reasons, diarrheal diseases represent a long-standing and significant burden, particularly in Latin America, Southeast Asia, and sub-Saharan Africa (6). Diarrheal diseases are caused by a range of pathogens, such as bacteria, viruses, and parasites including the *Entamoeba* genus. *E. histolytica* is an extracellular parasite that causes human infection with variable outcomes ranging from asymptomatic colonization to diarrhea, invasive colitis, liver abscess, and metastatic infection. Amebiasis from *E. histolytica* is the second leading parasitic cause of death globally but remains classified as a neglected disease (7). Globally 50 million cases are reported per year, resulting in 2.2 million disability-adjusted life years and 55-100 thousand deaths annually (reviewed in (8)).

*Entamoeba* infection starts with the ingestion of the cyst stage from contaminated food or water. Presumably in the small intestine, the *Entamoeba* cyst molts from its chitinous shell and differentiates into the metabolically active trophozoite form through a process known as excystation. Trophozoites then attach to the intestinal epithelium where they undergo asexual reproduction and encystation. The infectious fecal cysts are shed and contaminate the environment to complete the parasite life cycle. Trophozoites can either stay contained within the intestinal tract or may disseminate to soft tissue organs like the liver, the lungs, or the brain (9–12), although in ~90% of the cases, the infection remains in the intestinal lumen (13, 14). The current murine intestinal infection model surgically delivers trophozoites into the cecum of animals and has provided immense insight into host-pathogen interactions but produces no cysts (15). Thus, modeling the critical developmental stage interconversion that *E. histolytica* undergoes between ingestion and colonization of the cecum is not yet possible (16). A model that includes the developmental changes of excystation and encystation would allow the field to understand the transmission of the pathogen and find targets for intervention (reviewed in (17)), as only viable parasites completely encysted and shed via the fecal-oral route can contaminate food and water and infect a new host.

The parasitology field has used species that naturally colonize animals to expand the knowledge of infectious diseases that are fastidious to culture or model in the laboratory. Murine pathogens like *Plasmodium chabaudi* have been pivotal to studying the *in vivo* pathology of malaria. For the *Entamoeba* field, *Entamoeba invadens*, a reptile-specific pathogen, has provided critical insights related to developmental changes like encystation. Here, we screened for a natural murine *Entamoeba* colonizer and developed an oral infection model using Swiss Webster (SW) mice. SW mice treated with paromomycin showed no *E. muris* colonization. We further determine infection location within the intestinal tract and excystation cues for the purified fecal cysts.

## Results

### C57BL/6 mice, but not Swiss Webster mice, are chronically colonized with a naturally occurring *Entamoeba* organism

We screened transgenic and wild-type animals within our vivarium facility (Fig 1A) using a Pan-*Entamoeba* PCR (Fig S1). We screened fecal material from males and females with ages ranging from newly weaned to mice that spent up to a 1 year in our facility. Eighty percent of the mice from the C57BL/6 background were colonized with an *Entamoeba* organism. To address if an *Entamoeba* is naturally occurring in other vivariums, we requested fecal samples from C57BL/6 mice from five collaborators around the country (n=18). We detected *Entamoeba* using a single-step PCR in samples from one other institution while in another we only detected the parasite using a 2-step, nested PCR. However, colonization was by no means ubiquitous as the majority of these institutions were PCR negative even when using the nested approach (Table S1). In contrast, all the SW mice from our vivarium tested were PCR negative, regardless of age or sex (Fig 1A).

**Fig 1.**
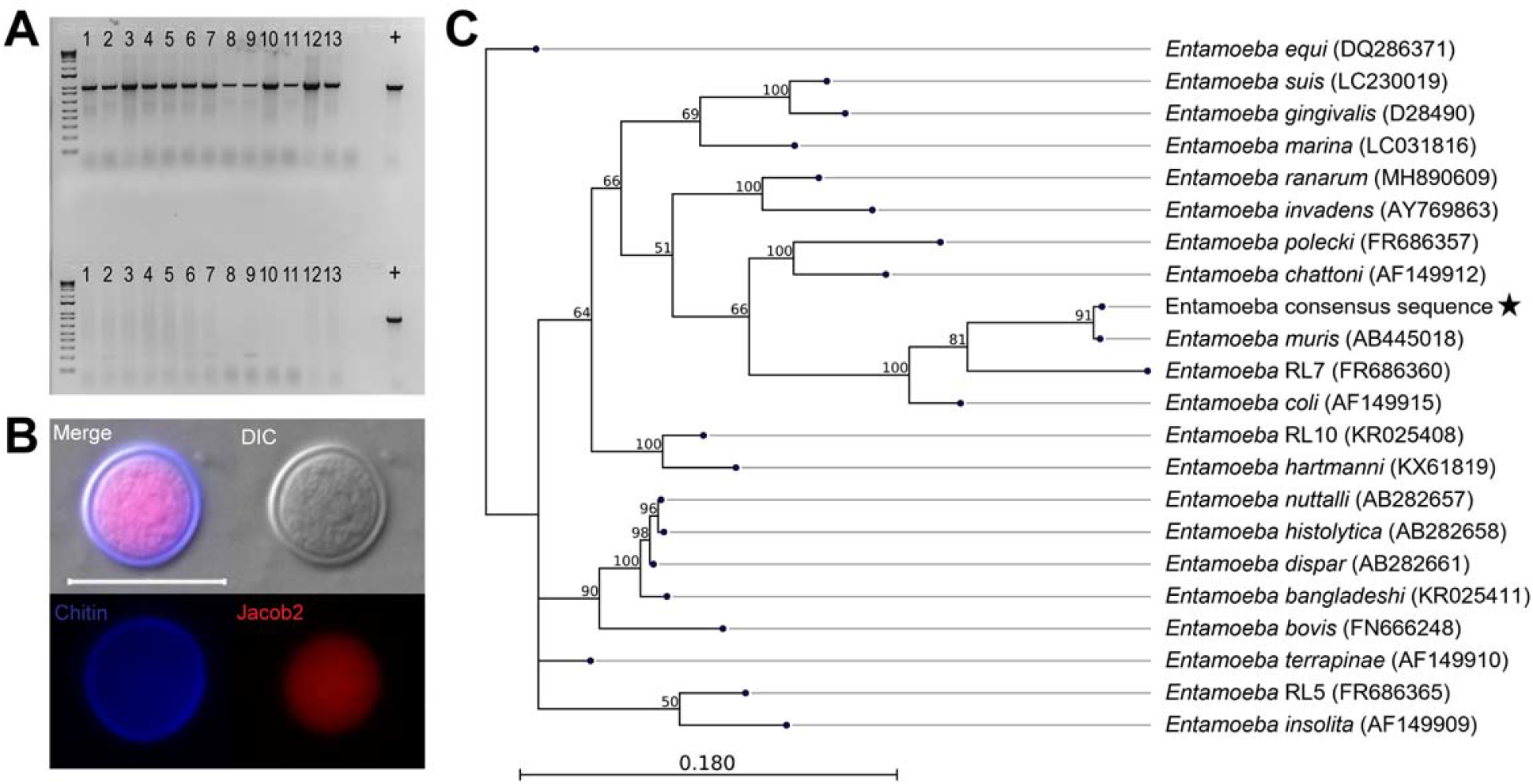
Identification of a murine *Entamoeba* species. (A) Representative Pan-*Entamoeba* PCR for C57BL/6 mice (top) and Swiss Webster mice (bottom) within our vivarium. The loading control was murine GAPDH (data not shown). Each lane number represents a mouse cage. Positive control (+) is isolated genomic DNA from axenic *Entamoeba histolytica* culture. (B) Immunofluorescence assay staining for chitin (Calcofluor White), Jacob2 (1A4 antibody), scale bar represents 20 μm. (C) 18S phylogeny is based on a 1,033 bp alignment including 27 published *Entamoeba* sequences, labeled as species name (NCBI accession), plus the *Entamoeba* found in our vivarium, indicated by a star. Numbers on branches are bootstrap values (%) based on 1,000 replicates (values >50% are shown). The scale bar indicates nucleotide substitutions per site.

To confirm the PCR findings in our animals, we developed a sucrose density gradient protocol based on fecal isolation methods from the *Entamoeba* literature (18) (Fig S2) and conducted phenotypic characterization of the isolated cysts (Fig S3). We processed feces of PCR-positive mice within a sucrose gradient of 1.33 specific gravity and isolated structures of 15 - 20μm in diameter. These structures were cyst-like and stained with calcofluor white, indicating the presence of chitin (Fig 1B). We further characterized these cysts by immunofluorescence detection of a previously published *Entamoeba-specific* antibody targeting the lectin Jacob2 (19). The sucrose gradient results for B6 and SW mice were 100% replicative of the Pan-*Entamoeba* PCR; SW mice did not display cyst-like structures in their fecal samples, while the B6 fecal samples contained 15-20 μm-diameter cysts that stained positive for both chitin and Jacob2 (Fig 1B).

To determine the species of *Entamoeba* present in our mice, we gel-purified and Sanger-sequenced the 1 kb pan-*Entamoeba* PCR product from cecum, colon, and fecal samples from 8 mice (n=24 total). The resulting reads were 100% identical to each other and most closely matched *Entamoeba muris* (GenBank accession number AB445018) with 98% query coverage at 92% identity and an E value of 0.00. We then performed phylogenetic analysis to further confirm this preliminary species identification. Alignment of 21 published *Entamoeba* 18S sequences from NCBI GenBank along with our consensus Sanger sequence yielded a final alignment length of 1,033 bp. A maximum-likelihood phylogeny built from this alignment shows our organism to cluster with *E. muris*, as expected from the BLASTn results, and form a clade with *Entamoeba* RL7 and *E. coli* (Fig 1C). Pairwise comparison of the *E. muris/coli* clade based on the 18S phylogenetic tree across the entire length of the alignment shows our organism to be 91.63% identical to *Entamoeba muris* with 9 total alignment gaps (0.87%) as compared to 81.82% identity and 42 gaps (4.1%) with its next-closest relative, *Entamoeba* RL7 (Table S2). Thus, we will refer to this organism as *E. muris* hereafter.

### Swiss Webster mice are susceptible to *Entamoeba muris* oral challenge

As SW mice were not currently colonized, we determined if these mice were suitable for *E. muris* infection and characterization using an oral challenge model. When orally challenged with a low cyst dosage (1×10^5^ isolated from B6 mice), SW mice were able to host a patent infection, as evidenced by fecal cyst shedding, but displayed asymptomatic infections in all cases. We monitored changes in weight during the infection but observed no significant differences between uninfected animals and animals orally challenged with *E. muris* cysts, although there was a trend for infected animals to have less weight gain when compared to uninfected controls (Fig 2A). Weight loss was only observed for one of the biological replicates (Fig S4A) and was not correlated to cyst shedding (Fig S4B). Fecal samples showed cysts as early as 3 days post-infection (dpi) via sucrose gradient. When using this method, we determined the peak of infection to be 7 dpi based on average cysts counts across four independent infection replicates (n=27 mice, 17 infected and 10 uninfected controls). Shedding per biological replicates does not show significant differences between biological replicates (Fig S4B). We detected a significant decrease in cyst shedding by 11 dpi, and very few *E. muris* cysts were detected using this method by 28 dpi.

**Fig 2.**
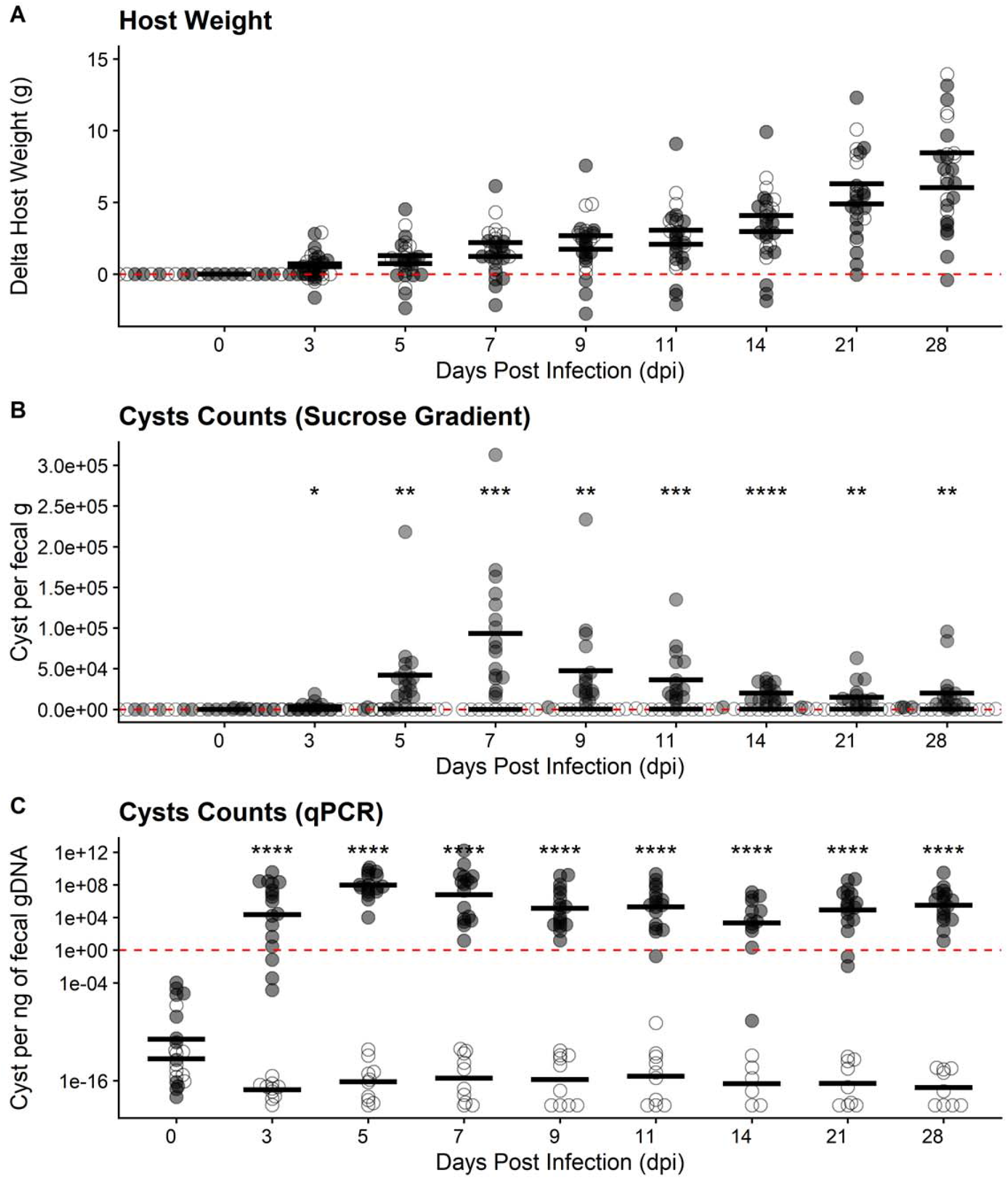
Swiss Webster mice are susceptible to *Entamoeba muris* oral challenge. (A) Host weight was monitored through the course of infection. (B) Quantification of cysts isolated by sucrose gradient from Swiss Webster fecal samples (normalized by fecal mass). Peak of infection was determined to be 7 dpi. (C) Quantification of cysts in fecal samples via qPCR isolated from Swiss Webster’s fecal samples (normalized by gDNA per qPCR reaction). Each dot represents a single mouse (n=27 mice, 17 infected and 10 uninfected controls). Open circles represent uninfected mice while gray circles represent infected mice. Significance was determined using a two-tailed t-test between the uninfected vs. infected average per DPI. Data combines four independent biological replicates (see supplement Fig S4 for individual biological replicate (n=4) plotting).

Using a similar approach to the Pan-*Entamoeba* screen, we designed qPCR detection primers that amplified a 200-bp amplicon to quantify *E. muris* shedding. We generated a standard curve using cyst samples of known concentrations based on counts, ranging from 1,562 cysts to 100,000 cysts on a 2-fold scale (Fig S5A). Our qPCR results determined that *E. muris* was detectable as early as 3 dpi, in accordance with the sucrose gradient isolation. However, the peak of infection was at 5 dpi, and a significant reduction was evident by 9 dpi. Taken together, these results suggest that qPCR detection occurs prior to peak viable cyst isolation.

### Paromomycin treatment prevents colonization of *Entamoeba muris*

Patients infected with *Entamoeba* species are prescribed antibiotics depending on the degree of pathogenicity. Two common drug treatments that are prescribed to patients infected with an *Entamoeba* species are metronidazole and paromomycin. For invasive disease, metronidazole is the gold standard for treatment, but some *Entamoeba histolytica* strains can develop resistance over time (20). Paromomycin is the treatment of choice for infections contained to the gastrointestinal tract due to its specificity against anaerobic microbes, but it is often prescribed in conjunction or soon after metronidazole treatment. To determine if paromomycin recapitulates its amoebicidal effect against *E. muris*, we orally infected SW mice with 7×10^4^ cysts isolated from B6 mice and immediately began treatment with paromomycin (16 g/L) in their drinking water for the first 7 days post-infection before reverting to untreated drinking water. Using the approaches presented earlier, we monitored *E. muris* colonization and host weight for 2 weeks post-infection. Overall, SW mice treated with paromomycin showed higher variability in weight changes throughout the course of treatment when compared to untreated mice (Fig S6). Infected mice treated with paromomycin trended toward slower weight gain than their uninfected treated counterparts, although this observation was not statistically significant (Fig S6). In contrast, both uninfected and infected mice given untreated drinking water displayed similar weight changes over the course of 14 days (Fig S6). As expected, SW mice that were challenged with *E. muris* but received no paromomycin treatment shed fecal cysts as early as 3-5 dpi via sucrose gradient (Fig 3A), even at a lower infectious dose of 7×10^4^ cysts. In concordance with our previous results, cyst shedding peaked at 7 dpi and declined significantly by 9 dpi (Fig 3A). In contrast, SW mice challenged with *E. muris* and treated with paromomycin never shed any viable *E. muris* throughout 14 dpi as determined by sucrose gradient isolation (Fig 3A).

**Fig 3.**
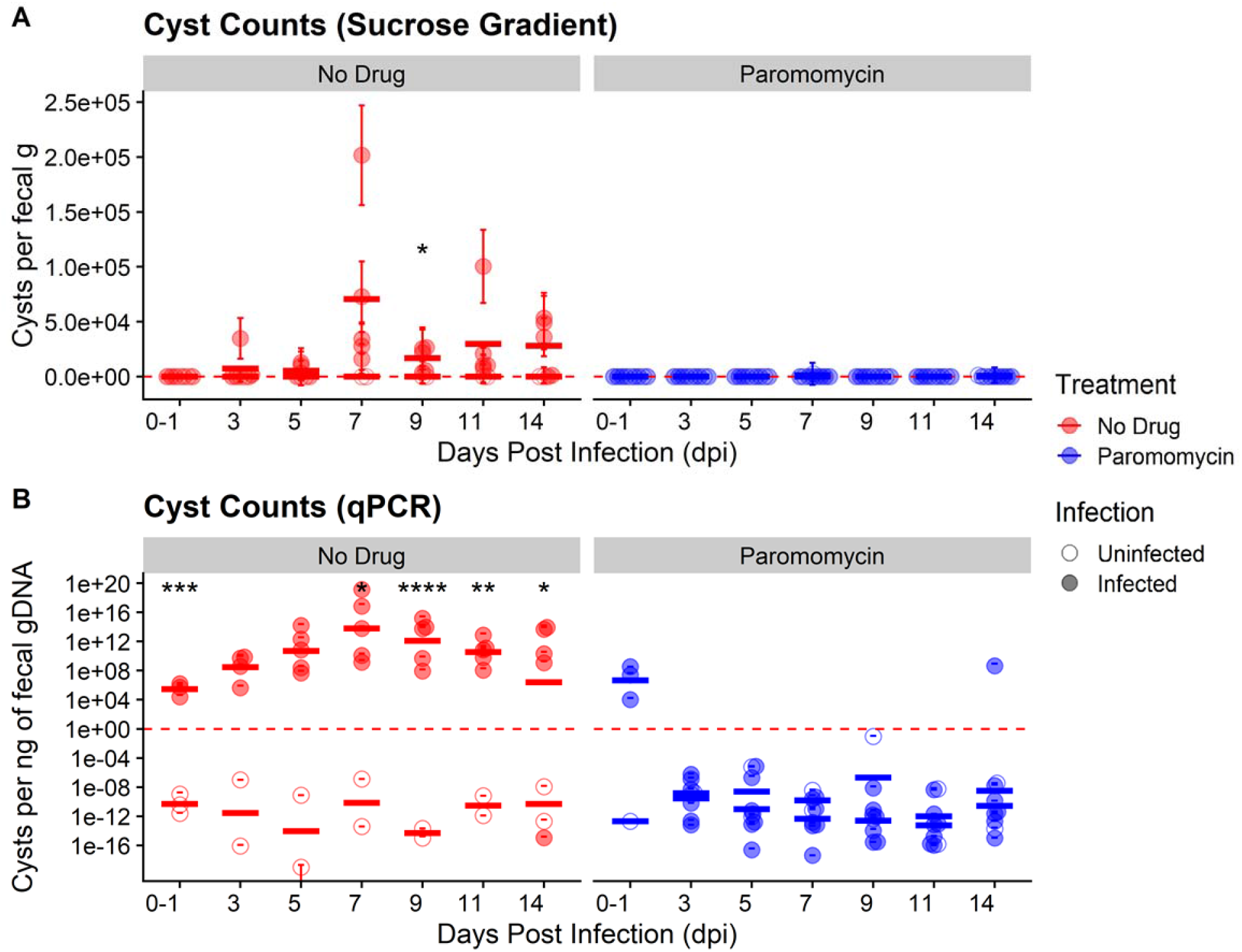
Paromomycin effectively inhibits colonization of SW mice by *Entamoeba muris*. (A) Quantification of cysts isolated by sucrose gradient from Swiss Webster fecal samples (normalized by fecal mass). (B) Quantification of cysts in fecal samples via qPCR isolated from Swiss Webster fecal samples (normalized by gDNA per qPCR reaction). Each circle represents a single mouse (n=16 mice, 12 infected and 4 uninfected controls). Open circles represent uninfected mice while filled circles represent infected mice. Red circles represent untreated mice while blue circles represent mice treated with paromomycin. Bars indicate calculated mean values for each experimental group per DPI. Significance was determined using a two-tailed t-test between the uninfected vs. infected average per DPI. Data combines two independent biological replicates.

We also monitored *E. muris* shedding via qPCR as a second method of detection. In agreement with our sucrose gradient data (Fig 3A), the peak of infection in untreated mice challenged with *E. muris* was also found to be 7 dpi by qPCR (Fig 3B). Unlike our sucrose gradient data, qPCR results indicated that all infected mice did have detectable *E. muris* between the day of infection (0 dpi) to 1 dpi (Fig 3B). However, infected mice treated with paromomycin no longer shed detectable *E. muris* by 3 dpi (Fig 3B). These mice continued to lack detectable *E. muris* up to 14 dpi, except for one mouse on day 14 (Fig 3B). Taken together, these results suggest that paromomycin effectively inhibits *E. muris* colonization when administered as a prophylactic and that our model can be used for drug screening studies relevant to luminal *Entamoeba* infections.

### *Entamoeba muris* resides in the large intestine at 5 days post-infection

*E. histolytica* is thought to replicate in the colon and has been found during diagnostic colonoscopies (21). As our model uses the natural oral route of infection, we aimed to determine where *E. muris* is located during primary infection. We infected SW mice with 10^5^ cysts by oral gavage and collected the intestinal content and mucus layer of murine gastrointestinal sections at 5 dpi, when most of the animals were shedding cysts by qPCR detection (Fig 2C). The small intestine was sectioned into three parts: duodenum (D), jejunum (J), and ileum (I). The cecum (Ce) and the colon (Co) correspond to the large intestine (Fig 4A). As a positive control for the presence of cysts, we included two fresh fecal pellets (F) and we used a standard loading control, murine GAPDH (Fig 4B, lower panel). As expected from clinical data (reviewed in (22)), *Entamoeba* localizes within the large intestine (Fig 4B, upper panel). While we found mouse to mouse variability between levels of *E. muris* detection within the cecum, colon, and fecal samples, no *Entamoeba muris* gDNA was isolated from the small intestine (Fig 4B, upper panel).

**Fig 4.**
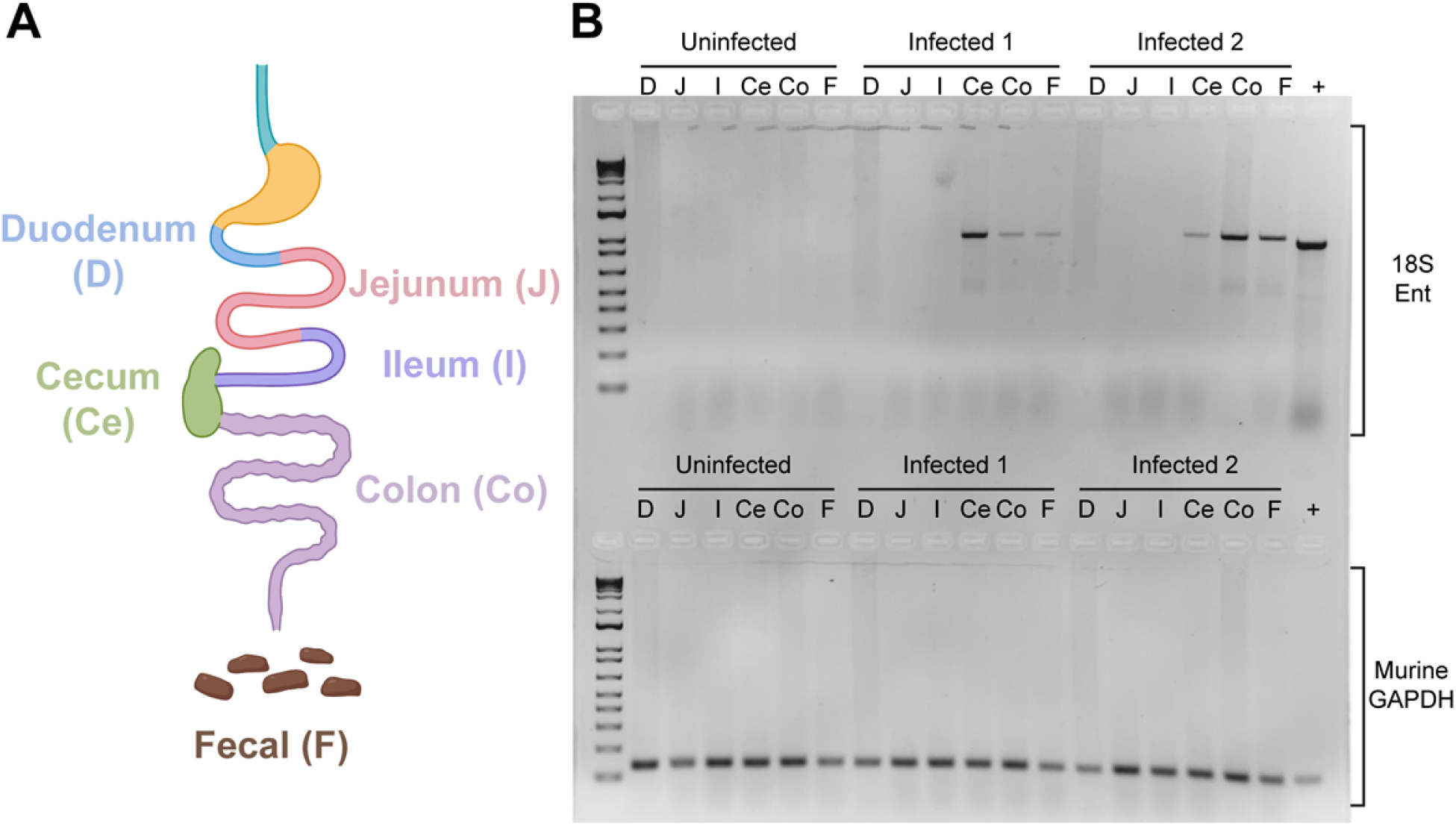
*Entamoeba muris* localizes to the large intestine of infected animals. (A) Schematics of the murine intestine. (B) The *Entamoeba* 18S gene was amplified from gastrointestinal sections (gel top). The positive control is genomic DNA extracted from an *Entamoeba histolytica* axenic culture. As a loading control, the murine GAPDH gene was amplified from various gastrointestinal sections, with gDNA isolated from mice tail snips serving as a positive control (gel bottom). Gel image is a representative of two independent biological replicates (n=6, 4 infected and 2 uninfected controls).

### Bile extract triggers *Entamoeba muris* excystation *in vitro*

To determine if the previously published *E. invadens* cues could trigger consistent *E. muris* excystation *in vitro*, we incubated the isolated cysts with Nanopure water, 80 mM sodium bicarbonate, 1% bovine bile, or a combination of both treatments for 24 hours (23). We scored excystation efficiency based on the percentage of the parasite that was outside of the chitin shell. An intact cyst was given a score of 0 (Fig 1B). We scored an open cyst with less than 50% of the trophozoite-mass excysted as a 1, cysts where 50% or more of the parasite was outside of the chitin shell as a 2, and empty chitin shells were given a score of 3 (Fig 5A). We observed excystation to be an asynchronous process, as scores ranged within each condition (Fig S7). Treatment of cysts with only 1% bovine bile resulted in greater than 70% excystation by 24 hours, which was statistically higher than the excystation rate of the Nanopure water treatment (*p* = 0.0040). This excystation rate was not enhanced by the addition of the sodium bicarbonate, and sodium bicarbonate alone did not significantly enhance excystation compared to water only (Fig 5B). These results strongly indicate that 1% bile is sufficient to trigger excystation, which implies that excystation of *E. muris* is occurring in the small intestinal tract as we would expect.

**Fig 5.**
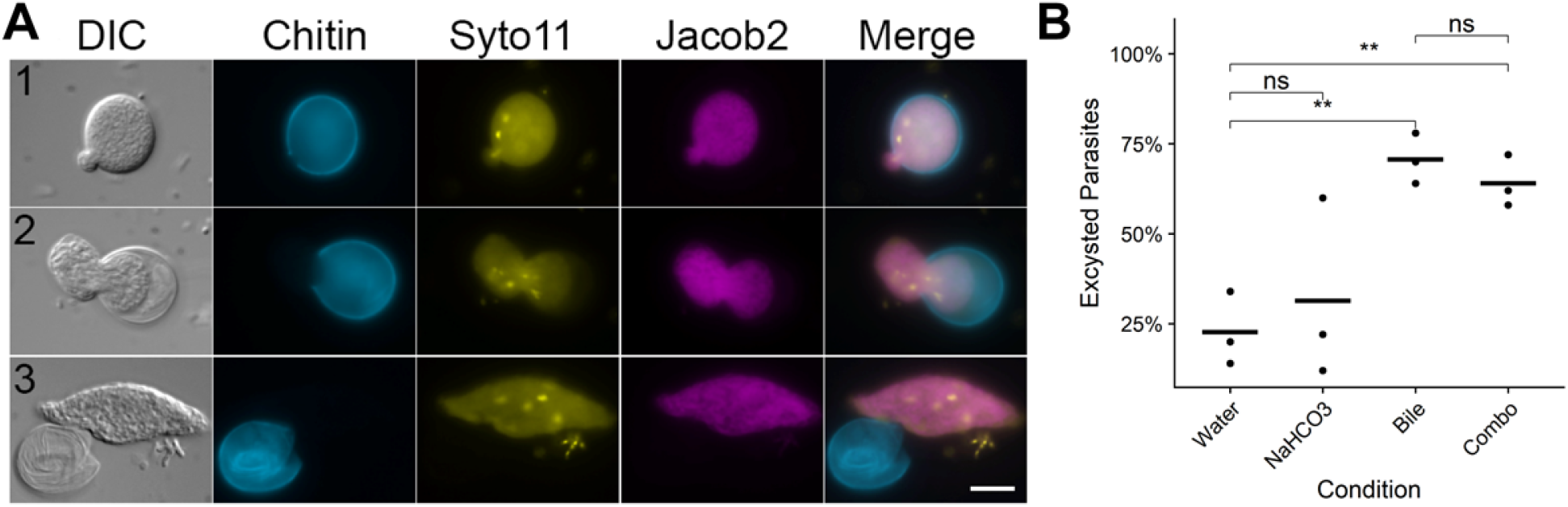
*Entamoeba muris* shows reliable excystation *in vitro* when treated with upper gastrointestinal tract components. Fecal cysts were purified by sucrose density gradient and then acid washed (0.1 M HCl). Cysts were inoculated into excystation conditions (1% bovine bile, 80 mM sodium bicarbonate, or a combination of both), then incubated for 24 hours at room temperature. (A) Cysts were scored from 0 to 3, where 0 represented an intact cyst and 3 is an empty chitin shell. Chitin (Calcofluor White), Jacob2 (1A4 antibody (17)), and nuclei (Syto11). Scale bar represents 10 μm. (B) Excystation rates (score >= 1) were quantified in these conditions. Significance was determined using a two-tailed t-test. Only significant pairwise comparisons are shown; Bile (*p*=0.004) and, NaHCO_3_ + Bile (*p*=0.0064). Each dot represents a biological replicate (n=3 independent experiments), black horizontal line is the average of the three biological replicates.

## Discussion

To the best of our knowledge, this work is the first to demonstrate that C57BL/6 mice can be chronically colonized with *E. muris*. C57BL/6 mice have been previously described as naturally resistant to *E. histolytica* when injecting trophozoites directly into a surgically exposed cecum (24). However, other protozoans like *Tritrichomonas musculis* have been reported to chronically colonize many colonies on the East Coast of the United States (25). *T. musculis* was found to change the immune response to protect against pathogens in mice with chronic colonization, and *E. muris* may also be changing the immune responses. Understudied intestinal protozoans may account for the variability of results between research institutions. Many of the fecal samples from C57BL/6 mice across the US did not amplify *Entamoeba*, but this lack of amplification is not due to low gDNA isolated from the feces, as we can still detect host gDNA in *Entamoeba* negative samples when using a primer set for murine GAPDH. One of the limitations of this study is that the *Entamoeba* identification is based on the 18S gene (Fig 1C and Table S2) and phenotypic characterization of the number of nuclei (>4) via microscopy (Fig S3); thus, further characterization might be required.

One surprising result was the positive Jacob2 staining for the *E. muris* cysts. The 1A4 antibody was previously described to distinguish *E. histolytica* cysts without cross-reacting either to *E. dispar* or *E. bangladeshi* cysts. The 1A4 antibody was generated against the flexible, serine-rich spacer of the Jacob2 lectin in *E. histolytica* (19), so perhaps this region is similar in *E. muris*. It is also interesting that the *E. Muris* cyst Jacob2 staining is associated with the pre-trophozoite during excystation and not the chitin-rich wall (Fig 4), as previously shown in stool samples and xenic cultures with another anti-Jacob2 antibody (26).

We demonstrated that previously uninfected SW mice can be infected with *E. muris* via oral gavage. We were especially interested in the natural route of infection as it has been established that the route of infection impacts disease progression for oral parasites like *Toxoplasma gondii* (27, 28). Because SW are outbred mice, they have been used for the evaluation of vaccines due to their unbiased immune response (29). In contrast, inbred mice of various genetic backgrounds exhibit different immune responses to an infectious challenge (30), specifically for parasitic infections that are intracellular (31) or extracellular (32). Inbreeding within human populations has been linked to protection against malaria (33) but inbreeding in wild European badgers intensified sex- and age-dependent tuberculosis disease (34). Future studies will examine *E. muris* oral infection in other inbred and outbred mice as well as immune deletion strains to determine the inflammatory responses necessary for *Entamoeba* control.

We also demonstrate that SW mice can be protected from *E. muris* infection oral infection using paromomycin. A surprising result from these experiments was that mice treated with paromomycin exhibited higher variability in weight changes when compared to untreated mice (Fig S6). This observation may hint at subtle differences in how a host responds to antibiotic treatment. In addition, paromomycin served as an effective prophylactic by inhibiting *E. muris* colonization after oral infection, demonstrated by a complete lack of cyst shedding as quantified by sucrose gradient (Fig 3A) and no qPCR detection by 3 dpi (Fig 3B). However, most patients do not take anti-parasitic drugs as prophylaxis, but rather as treatment for an already established infection. In our paromomycin treatment studies, we did observe early detection of *E. muris* in infected mice by qPCR at 1 dpi but not by 3 dpi. This early but transient detection could be attributed to *E. muris* cysts passively shed in the feces as a byproduct of oral infection or early colonization of parasites that were later killed by paromomycin treatment. While more studies are needed to demonstrate that paromomycin can directly kill an active *E. muris* infection in SW mice, these experiments are a proof of principle that our *E. muris* oral infection model can be applied to both characterization of currently available anti-parasitic drugs and discovery of novel therapeutics against *Entamoeba* spp.

Contrary to our expectations, about 20% of cysts isolated from unfixed fecal material via a sucrose gradient undergo asynchronous excystation when stored overnight at 4°C in Nanopure water. These results are surprising because, for axenic *Entamoeba invadens*, a combination of cues encountered in the upper gastrointestinal tract are required for comparable levels of excystation (23). Perhaps chemical signals present in the fecal samples, not eliminated during the sucrose gradient purification and not present in *E. invadens* literature, are triggering excystation in the isolated cysts. It may also be possible that exposure to sucrose during density gradient purification may serve as a nutritional cue for *E. muris* to excyst at low levels. The experimental induction of excystation with the bile treatment alone was also surprising because for *E. invadens*, bile alone yields less than 40% excystation (23). We did not perform the water pre-treatment as described in the *E. invadens* protocol. Perhaps our isolation process (Fig S2) might act as a water pretreatment, given the excystation yield of treatment with bile alone is comparable to the combination treatment previously reported (23).

There was a significant difference in the number of parasites quantified when using qPCR detection versus sucrose gradient isolation. This result is an important limitation that might be explained by the nature of the two selected assays. The sucrose isolation protocol selects for healthy cysts with a specific gravity of 1.33. A parasite that is in the trophozoite state, currently excysting, or that has a suboptimal cyst wall would be lost during the density gradient protocol. Meanwhile, the qPCR assay detects parasites in any state regardless of viability. In addition, each cyst can contain more than 4 nuclei, further increasing the amount of detectable *E. muris* gDNA in fecal samples.

During this project, we also discovered the importance of humidity on cyst viability, which has been characterized previously in other diarrhea causing parasites (35). Vivarium records indicate that there are drastic differences in humidity between the winter (20%) and summer (50%). The number of cysts isolated that were “healthy” and presumably viable at the specific gravity of 1.33 (Fig S2B, pellet 3) was dramatically reduced during the winter months. When we examined the waste sections of the gradient, where the material of different density would be expected, we found many cysts with a desiccated appearance (Fig S2B, pellet 2). Thus, room humidity will be important for researchers to monitor as they develop this model in their own facilities. Humidity may also play an important role in the seasonality that is seen with increases in human *E. histolytica* infections (36–38).

We foresee our model to be useful in *Entamoeba* stage interconversion research as well as drug efficiency testing. We hope for improvement in the methodology of detection by targeting moieties present specifically in the cyst stage. Lastly, we have confidence that with further studies, by our and other groups, the establishment of a robust culturing protocol is attainable to study parasite-microbiome interactions *in vitro*. We are excited to present these results, which allow for a myriad of new research avenues focusing on parasite physiology and transmission.

## Materials and Methods

All mice were treated according to the guidelines established by the Institutional Animal Care and Use Committee (IACUC) of the University of Wisconsin School of Medicine and Public Health (protocol #M005217). The institution adheres to the regulations and guidelines set by the National Research Council.

### Screen for colonized mice

Fecal samples were collected from various institutions within the continental United States (Table S2), as well as our own vivarium facility. Genomic DNA (gDNA) was isolated following previously published protocols with the following modifications (39, 40): briefly, whole feces (~0.10 g) were placed in solvent-resistant screw-cap tubes containing 0.1 mm zirconia/silica beads (BioSpec Products 11079101z) and 1 large stainless steel bead (BioSpec Products 11079132ss) suspended in 20% SDS buffer (200 mM Tris·HCl, pH 8.0/200 mM NaCl/20 mM EDTA) and UltraPure Phenol/Chloroform/Isoamyl alcohol, pH 7.9, 25:24:1 (Invitrogen 15593-049). Samples were bead beat on high for 3 minutes at room temperature, and gDNA was precipitated with 3 M sodium acetate and isopropanol overnight. gDNA was cleaned using DNA Clean & Concentrator 5 (Zymo Research D4004). For identification of *Entamoeba muris*, a set of pan-*Entamoeba* primers were designed by downloading full-length *Entamoeba* 18S rRNA sequences (n=63, 25 *Entamoeba* species) from NCBI GenBank, aligning them in CLC Genomics Workbench v20.0.4 (Qiagen, Hilden, Germany), and identifying conserved regions to target forward and reverse primers (Forward: 5’-AGATACCGTCGTAGTCCT-3’ and Reverse: 5’-ACGACTTCTCCTTCCTCTAA-3’) which together amplify a 1 kb product (Fig S1, reaction 1). A total of 500 ng per PCR reaction were the genetic template for reaction 1, while for some samples a 2-step, nested PCR, was performed using 5 μl of reaction 1 as genetic template, using the same primer set and thermocycler conditions.

### Cyst Purification

#### Cyst counts

Fecal samples were processed used sucrose gradients as previously described with some modifications (41). Briefly, fecal samples (0.25–5 g) were ground to a fine powder using a mortar and pestle then shortly homogenized with Nanopure water for 15 minutes using a Mini Rotator (Glas-Col) at 60 rpm. The resulting solution was filtered through four-ply cotton gauze, and samples were pelleted for 10 minutes at 2500 x g. The resulting pellet layered on top of 1.5 M sucrose solution. The mid-layer was washed with Nanopure H_2_O and pelleted again at same speed. Isolated, un-fixed cysts were used as the input for oral infection.

### Immunofluorescence Assays

Fresh fecal sample was used to isolate cysts as described above then fixed in 10% formalin, washed twice, and resuspended. Isolated cysts were blocked for 5 minutes in 3% normal goat serum at room temperature with rotation. After washing, primary antibody 1A4 (19) was added at 1:1000 dilution (2.9 μg/ml concentration) and incubated for 2 hours with rotation. As the secondary antibody, we utilized a goat anti-mouse IgG conjugated to Alexa Fluor 594 (Thermo Fischer Scientific) and incubated in the same conditions overnight. A set of washes in between antibodies was conducted. Lastly, the samples were stained with 0.1% Calcofluor White Stain (Sigma-Aldrich) according to manufacturer instructions. To target nucleic acids, 0.025% Syto11 stain (Thermo Fisher Scientific) was used. Equal parts of sample and VECTASHIELD Mounting Media (Vector Laboratories) were utilized. Samples were visualized using an Axio Imager 2 microscope (Zeiss). Images were captured at 40X and 100X magnification using the DAPI, DIC, GFP and TexRed channels.

### Sanger sequencing and phylogenetic analysis

~1 kb products from the Pan-*Entamoeba* PCR above were gel purified using a Zymoclean Gel DNA Recovery Kit (Zymo Research) and submitted to the UW-Madison Biotechnology Center for Sanger Sequencing using the amplification primers described above. Sanger reads were manually inspected and edited using Sequencher v10.1 (Gene Codes Corporation) and queried against NCBI GenBank using Megablast (42) and default parameters. Twenty-seven full-length 18S *Entamoeba* sequences were downloaded from NCBI GenBank and aligned, along with our consensus Sanger sequence, using CLC Genomics workbench v20.0.4 (final length 1033 positions). A phylogenetic tree was inferred from the alignment with PhyML v.1.8.1 (43) using the general time reversible (GTR) substitution model and 1000 bootstrapped data sets were used to estimate statistical confidences of clades. To quantify nucleotide-level distances within the clade containing our organism, a pairwise distance matrix was constructed with the 4 clade members in CLC Genomics Workbench v20.0.4.

### Mouse Infections

#### Characterization of *E. muris* oral infection

House-bred male and female Swiss Webster Outbred mice were used to characterize *Entamoeba muris* infection for biological replicate 1. Male and female Swiss Webster (CFW) Outbred mice, purchased from Charles River Laboratories, were used to characterize *Entamoeba muris* infection for biological replicates 2-4 (Fig 2). Mice were 6-8 weeks of age at the time of oral challenge, individually caged, and provided enrichment for the duration of the experiments. All animals were gavage-fed either purified cysts (1×10^5^) or 1X PBS as a control.

#### Paromomycin treatment

Male and female Swiss Webster (CFW) Outbred mice were purchased from Charles River Laboratories and used to test paromomycin efficacy against *Entamoeba muris* oral challenge. Mice were 8-15 weeks of age at the time of oral challenge, individually caged, and provided enrichment for the duration of the experiments. All animals were gavage-fed either purified cysts (7×10^4^ due to limited input cyst amounts) or 1X PBS as a control. Treated mice were administered paromomycin sulfate (Research Products International) via drinking water (16 g/L) *ad libitum* for seven days before switching to normal drinking water. Throughout the length of the experiments, mice consumed an average of 5 mL per day with no difference in water consumption between untreated and treated mice.

#### Localization

House-bred male and female Swiss Webster Outbred mice were infected and euthanized at 5 dpi. The entire murine intestine was isolated and placed in 1X PBS. The small intestine was divided into three sections. Starting from the stomach, the first third was determined to be the duodenum, the following section was labeled as the jejunum, and the most proximal to the cecum was labeled ileum. For the large intestine, the entire cecum pouch and colon were used as independent sections. Intestinal contents of each section and a generous scraping of the host epithelial layer were pelleted at 2500 x g at room temperature for 5 minutes. The pellet was then processed in the previously described gDNA extraction protocol.

### Cyst Quantification

#### Sucrose Gradient

Cysts were purified as described above. For sucrose gradients, ~0.25 g fecal sample were used to isolate cysts. The cysts were then counted using a hemocytometer, and fecal mass was used to normalize counts.

#### qPCR

For quantification of *Entamoeba muris* by qPCR detection, standard curves were generated with known concentrations of cysts (FigS5) and intercalated dye (Bio-Rad SsoAdvanced Universal SYBR Green Supermix) using either a QuantStudio™ 7 Flex Real-Time PCR System or a StepOnePlus Real-Time PCR System (Applied Biosystems). Individual standard curves were generated specifically for each system. The standard curve for the QuantStudio 7 Flex PCR system ranged from 1,562–100,000 cysts (Fig S5A) and was used to calculate cyst concentrations in all biological replicates of Fig 2C and biological replicate 1 of Fig 3B. The standard curve for the StepOnePlus PCR system ranged from 1,562–50,000 cysts (Fig S5B) and was used to calculate cyst concentrations for biological replicate 2 of Fig 3B. Primers were designed as described above for pan-*Entamoeba* PCR (Fig S1), except here they were chosen to amplify a 200 bp product. The following primers were used: Forward: 5’-TCGAGATAAACGAGAGCGAAAG-3’ and Reverse: 5’-GTCAGGACTACGACGGTATCTA-3’. Fecal samples were collected, and ~0.10 g of whole feces were used as the starting material for gDNA isolation as described above. Per qPCR well, 100 ng of sample gDNA were loaded and analyzed. The total number of *E. muris* cysts present in each sample per nanogram of gDNA was calculated using the CT values of the experimental samples and the linear trendline equations of their respective standard curves.

### Excystation assay

Assays were conducted as previously described for *Entamoeba invadens* (23). Briefly, isolated cysts were acid washed with 0.1 M HCl for 10 minutes, followed by a second wash with Nanopure water. Cysts were then inoculated into each excystation treatment condition at a final amount of 10,000 cysts per condition: Nanopure water, 1% bovine bile, 80 mM sodium bicarbonate, or a combination of both bovine bile and sodium bicarbonate. Samples were incubated for 24 hours at room temperature, washed with Nanopure water, and fixed in 10% formalin. Fixed cysts were stained with 0.1% Calcofluor White. Cysts were mounted and visualized as described for immunofluorescence assays. A total of 50 cysts per biological replicate were scored per tested condition.

## Acknowledgments

We sincerely thank Upinder Singh and her lab for advice, and Jerry Cangelosi and his lab for the 1A4 anti-Jacob2 antibodies. We would like to thank researchers from across the US for providing mouse fecal samples. We also thank members of the Knoll lab (Nicole M. Davis, Billy J. Erazo Flores, Carlos J. Ramirez Flores, Katie M. Cataldo, and Jasmine N. Hughes) for assistance with experiments, software, as well as scientific advice. We thank Apoorva P. Maru and Sarah K. Wilson for their critical reading and editing of this manuscript.

---

**Fig S1.**
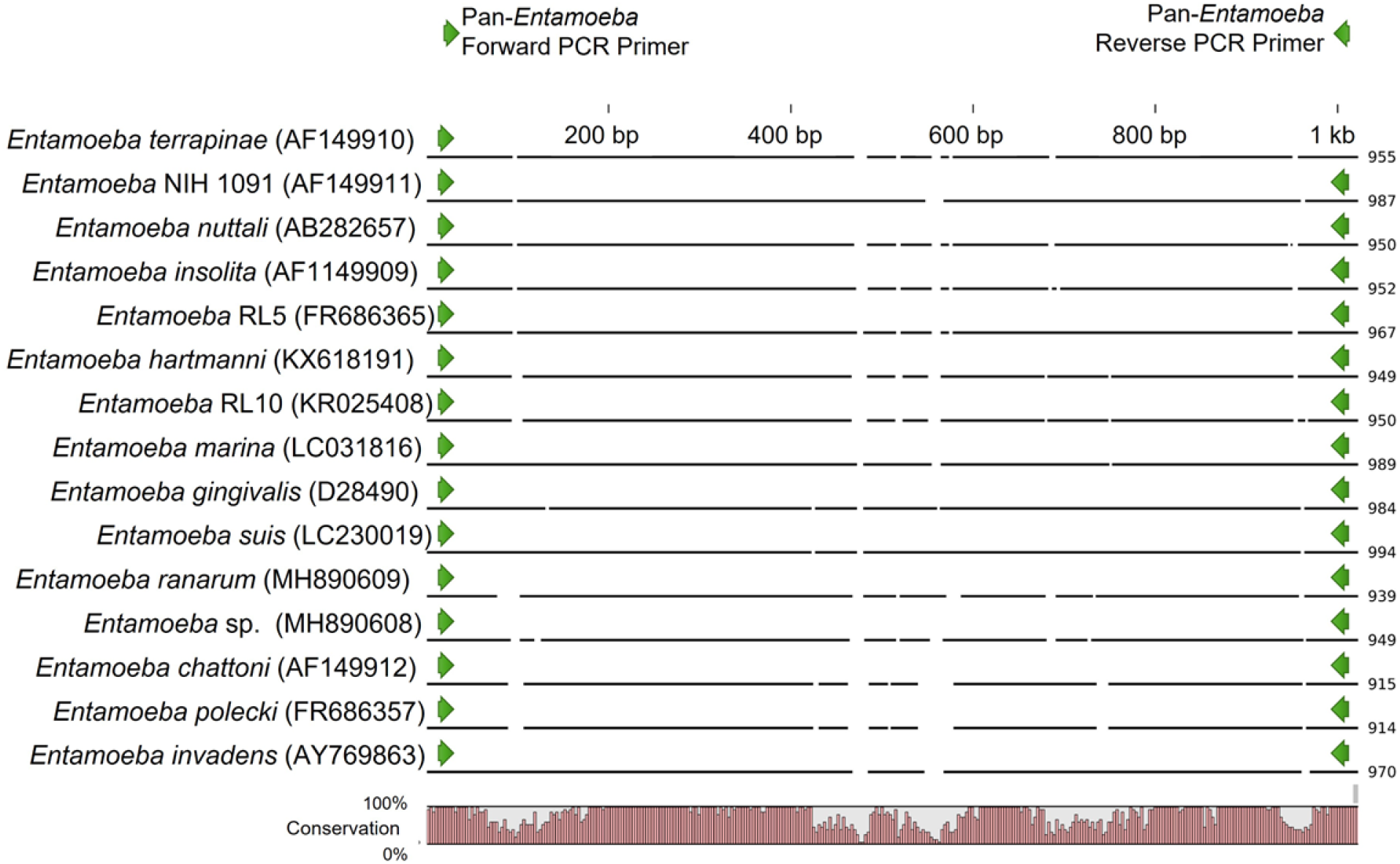
Pan-*Entamoeba* primer design. Full-length Entamoeba 18S rRNA sequences (n=63, 25 *Entamoeba* species) were downloaded from NCBI GenBank, aligned in CLC Genomics Workbench v20.0.4 (Qiagen, Hilden, Germany), and conserved regions were the target for the placement of the forward and reverse primers, which together amplify a 1 kb product. Forward: 5’-AGATACCGTCGTAGTCCT-3’ Reverse: 5’-ACGACTTCTCCTTCCTCTAA-3’

**Fig S2.**
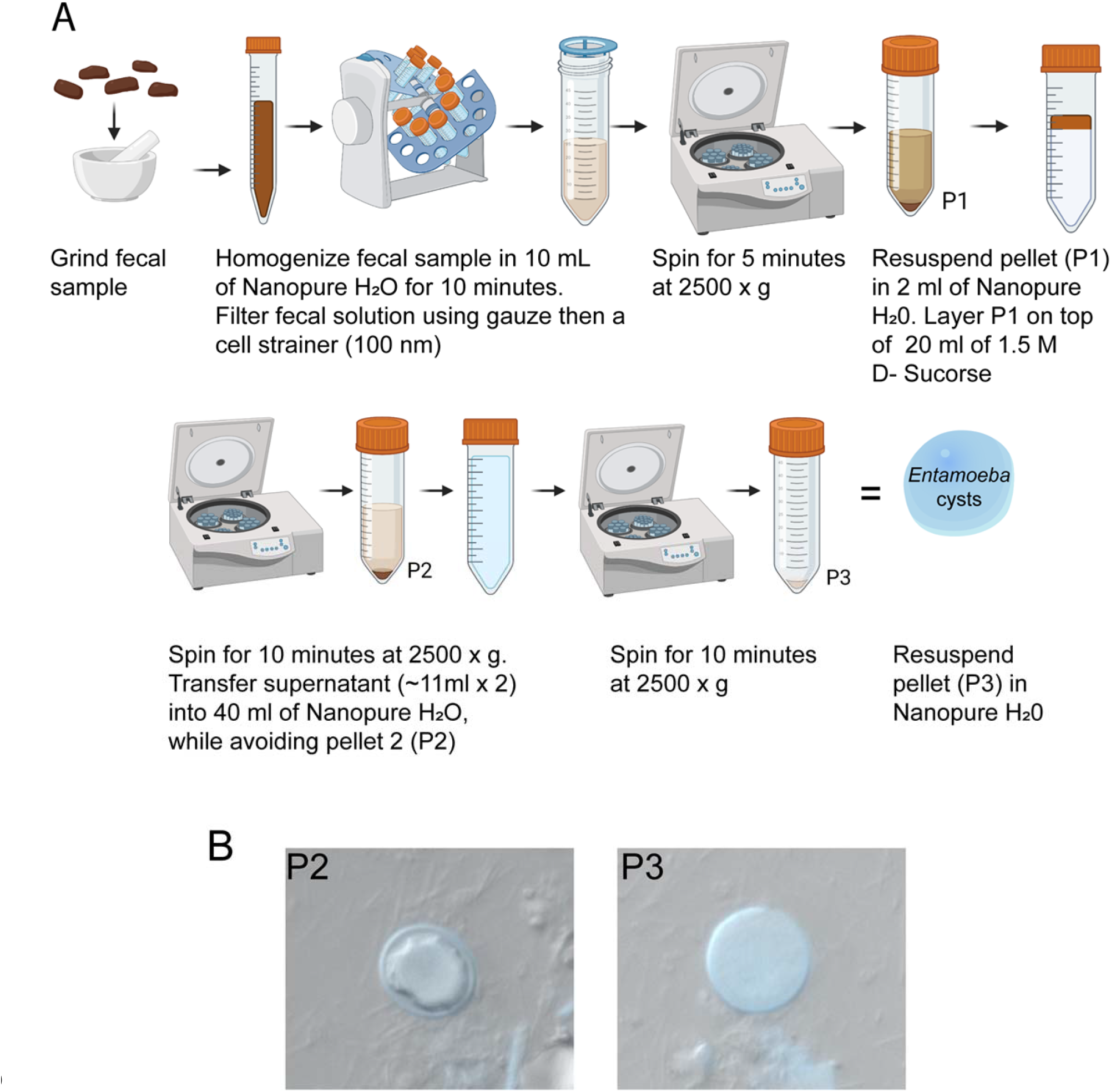
Sucrose gradient isolation protocol. (A) Fecal samples (0.25–5 g) that were collected overnight were processed as indicated above. Briefly, fecal samples were ground to a fine powder using a mortar and pestle then shortly homogenized with Nanopure water for 15 minutes using a Mini Rotator (Glas-Col) at 60 rpm. The resulting solution was filtered through four-ply cotton gauze, and samples were pelleted for 10 minutes at 2500 x g. The resulting pellet was layered on top of 1.5 M sucrose solution. The mid-layer was washed with Nanopure water and pelleted again at the same speed. Pellet 3 was suspended in Nanopure water, and the isolated un-fixed cysts were used as the input for oral infection. (B) While cysts can be found in the P2 pellet during the winter months, they have a dehydrated appearance compared to the cysts in the P3 pellet, likely due to the low humidity of the vivarium.

**Fig S3.**
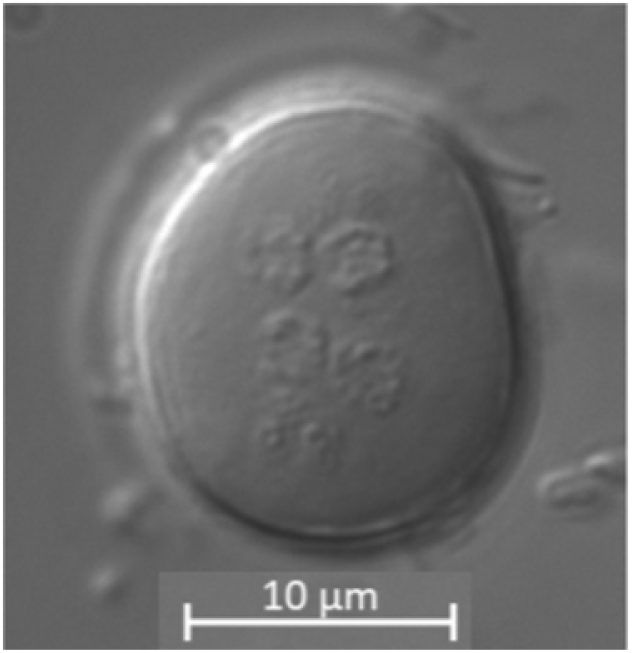
Representative phenotypic characterization of the number of nuclei of *E. muris* cysts (>4 nuclei). Image was taken using DIC. Scale bar represents 10 μm.

**Fig S4.**
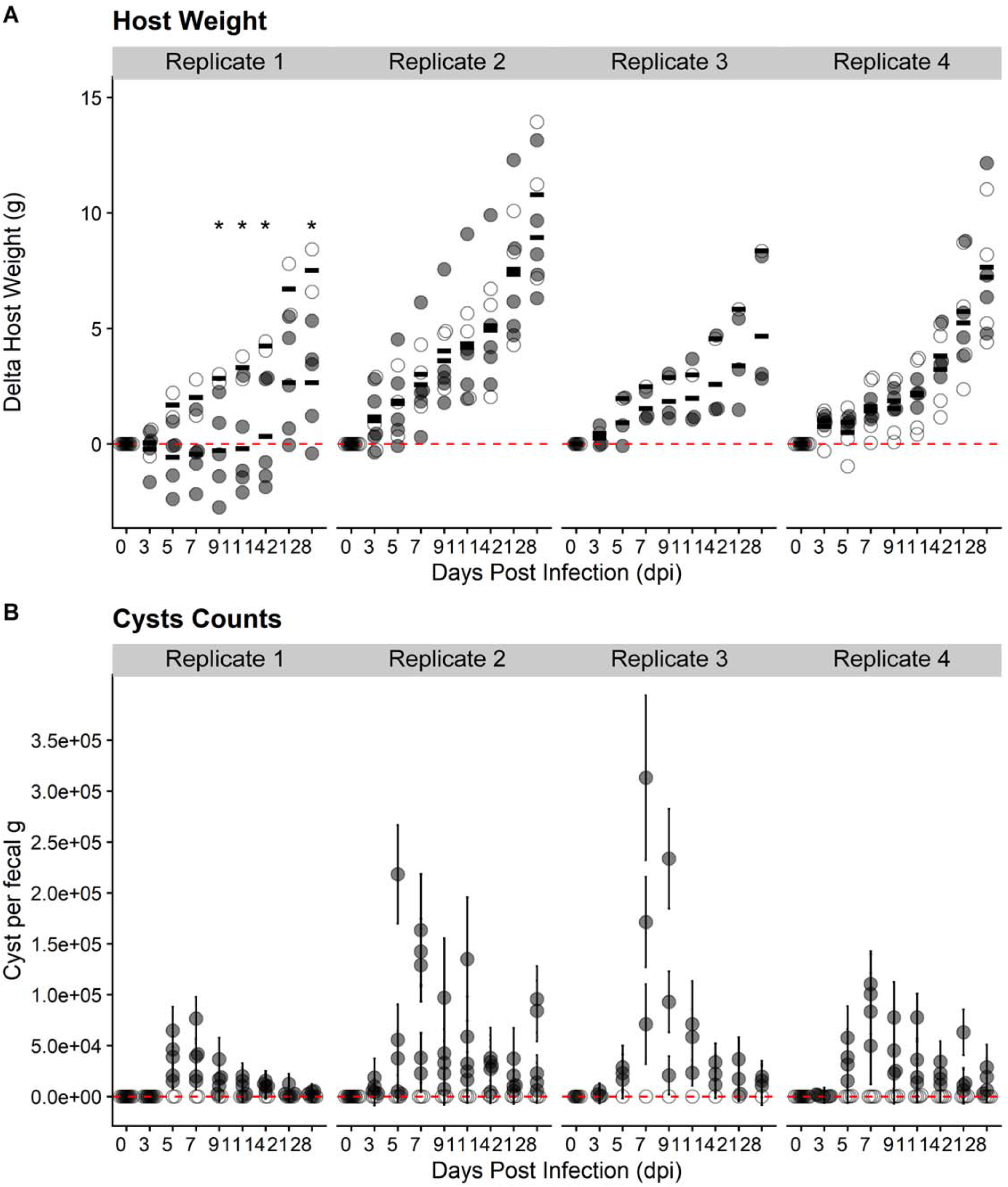
Swiss Webster mice are susceptible to *Entamoeba muris* oral challenge, per biological replicate. (A) Host weight was monitored through the course of infection. (B) Quantification of cysts isolated by sucrose gradient from Swiss Webster fecal samples (normalized by fecal mass). Each dot represents a single mouse. Open circles represent uninfected mice while gray circles represent infected mice.

**Fig S5.**
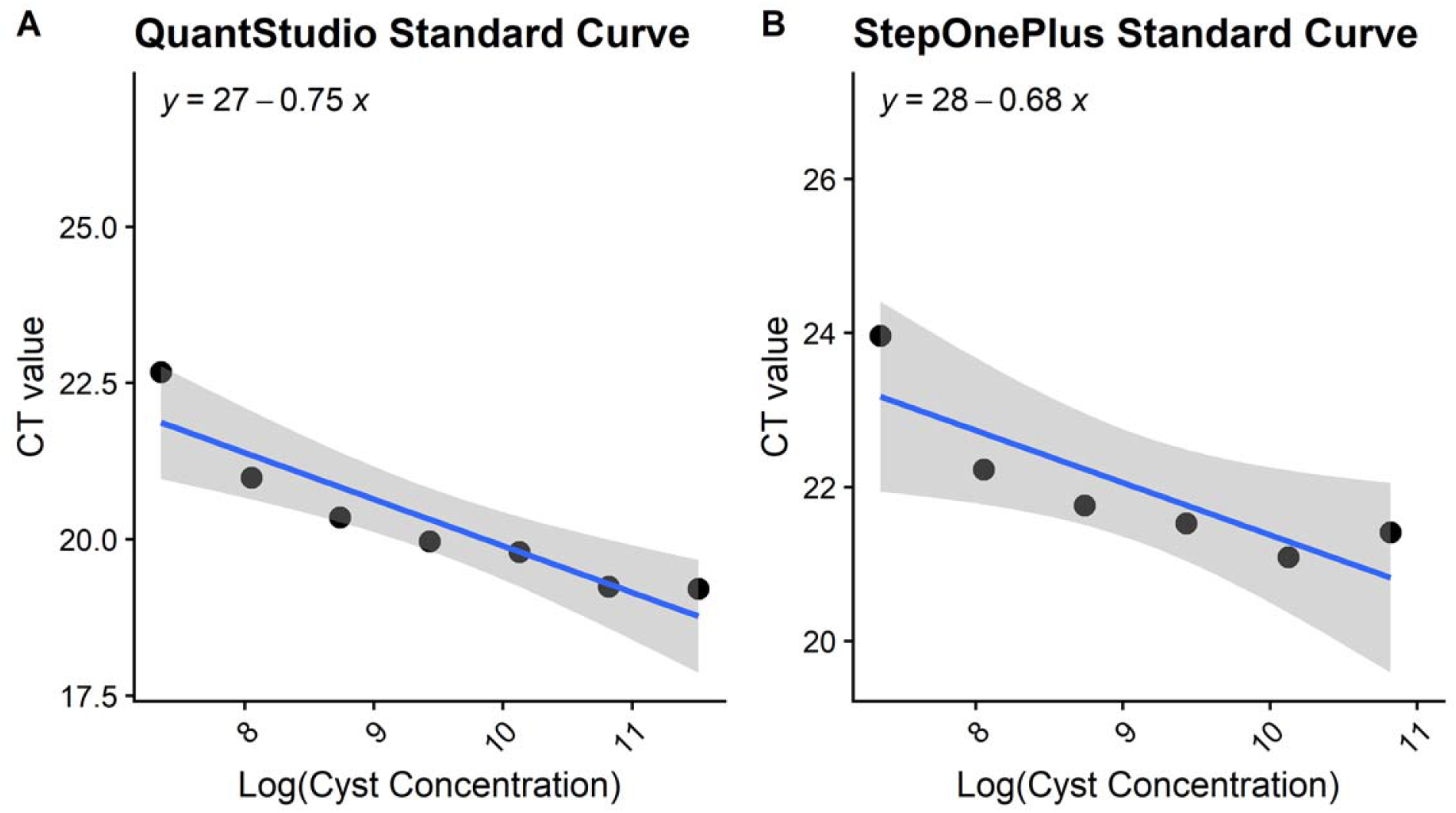
Standard curves for qPCR cyst quantification. Primers were designed as described above for pan-*Entamoeba* PCR (Fig S1), except here they were chosen to amplify a 200 bp product (see primers below). Two standard curves were generated for each real-time PCR system used in our studies using cyst samples of known concentrations, ranging from (A) 1,562 cysts to 100,000 cysts or (B) 1,562 cysts to 50,000 cysts expressed on a 2-fold scale. Concentrations for Figs 2C and 3B were calculated using the cycle threshold (CT) values of the experimental samples and the linear trendline equations presented above. Forward: 5’-TCGAGATAAACGAGAGCGAAAG-3’ Reverse: 5’-GTCAGGACTACGACGGTATCTA-3’

**Fig S6.**
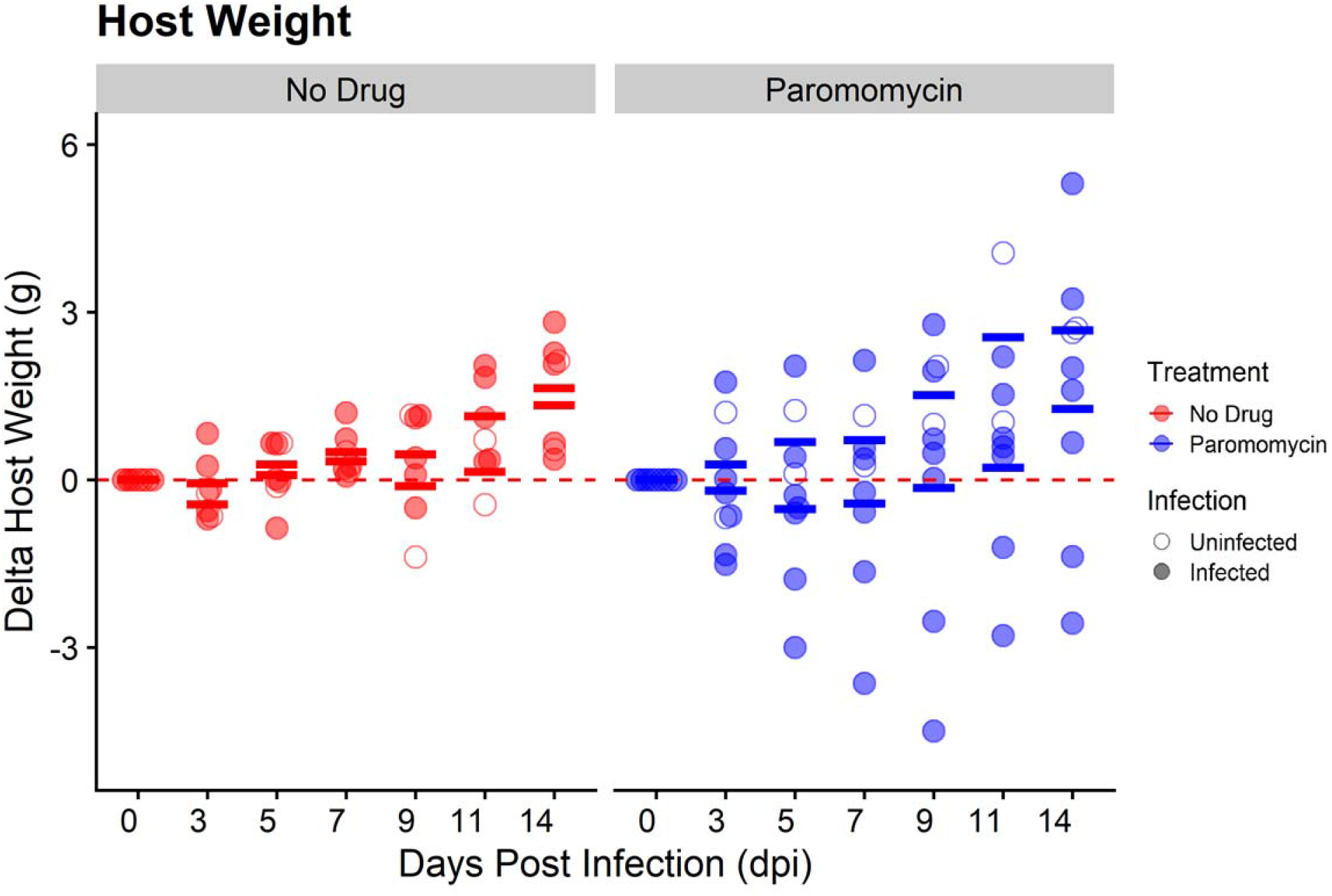
Paromomycin-treated SW mice display variable changes in host weight compared to untreated SW mice. (A) Host weight was monitored through the course of infection at 0, 3, 5, 7, 9, 11, and 14 dpi. Open circles represent uninfected mice while filled circles represent infected mice. Red circles represent untreated mice while blue circles represent mice treated with paromomycin. Bars indicate calculated mean values for each experimental group per DPI.

**Fig S7.**
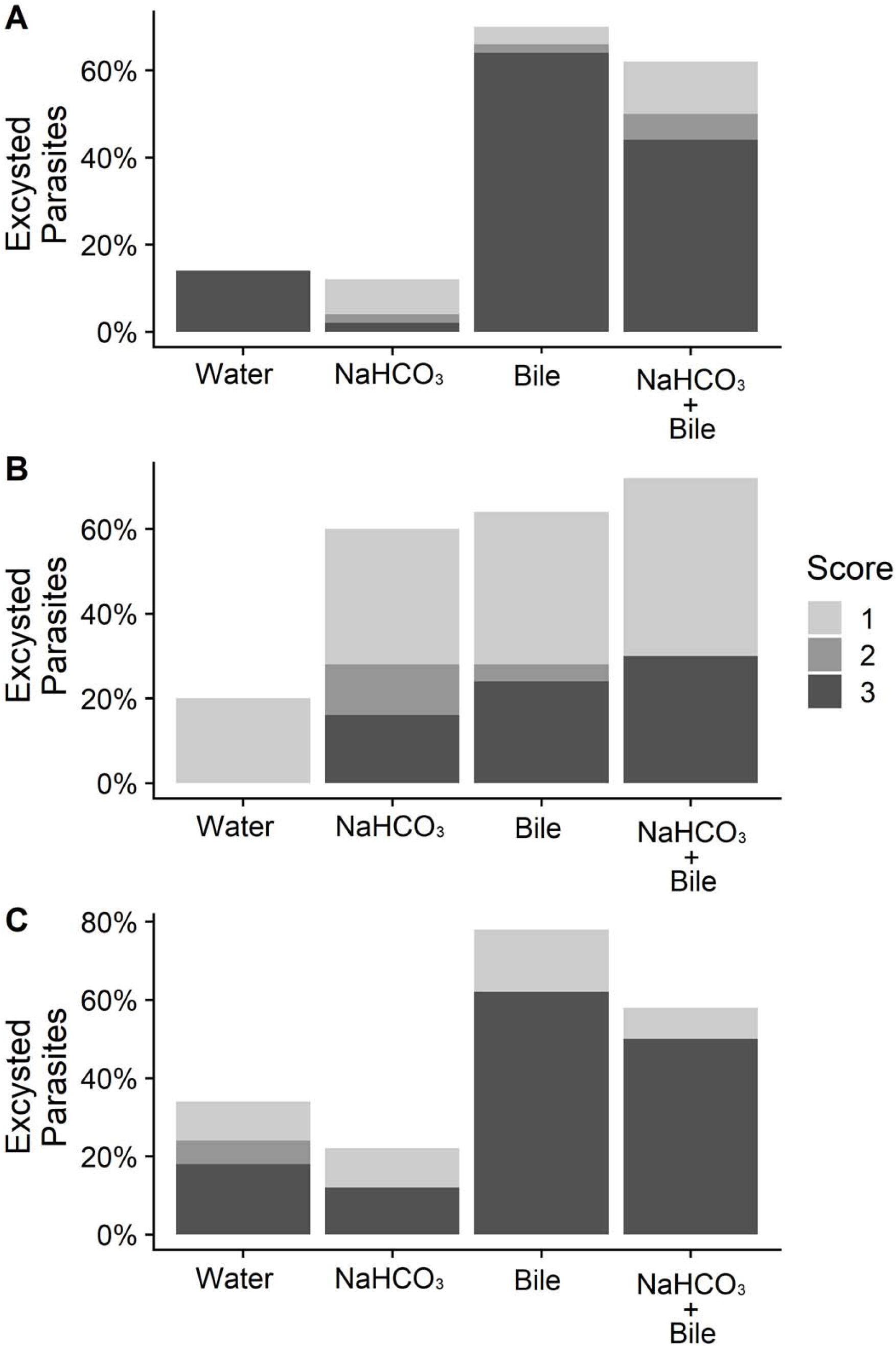
*Entamoeba muris* excystation is an asynchronous process. Fecal cysts were purified by sucrose density gradient and then acid washed (0.1 M HCl). Cysts were inoculated into excystation conditions (1 mg/ml bile, 80 mM sodium bicarbonate, or a combination of both), then incubated for 24 hours at room temperature. Cysts were scored from 0 to 3, where 0 represented an intact cyst and 3 is an empty chitin shell. Each panel (A-C) represents the individual biological replicates averaged in Fig 5.

**Table S1.** Colonization rates at vivarium facilities around the United States.

| Institution<br>Location | B6<br>Samples | % Positive<br>(40 cycle PCR) | % Positive<br>(Nested PCR) |
| --- | --- | --- | --- |
| Arizona | N= 4 | 0% | 25% |
| California | N= 2 | 100% | 100% |
| Pennsylvania | N= 2 | 0% | 0 % |
| Texas | N=1 | 0% | 0% |
| Vermont | N=10 | 0% | 0% |

**Table S2.** Pairwise comparison of *Entamoeba muris/coli* clade. Comparison is based on a 1,033 bp alignment. Percent nucleotide identity is shown above the diagonal and number of gaps are shown below the diagonal.

|  | *Entamoeba coli* | *Entamoeba RL7* | *Entamoeba muris* | Consensus sequence |
| --- | --- | --- | --- | --- |
| *Entamoeba coli* |  | 77.9 | 77.6 | 76.1 |
| *Entamoeba RL7* | 51 |  | 82.6 | 81.7 |
| *Entamoeba muris* | 44 | 37 |  | 91.6 |
| Consensus sequence | 49 | 42 | 9 |  |

## References

1. KD K, RG L, DD J, JM R, HC S. Intestinal parasitism in the United States: update on a continuing problem. The American journal of tropical medicine and hygiene. 1994;50(6).

2. R S, TH X, LNS H, MJ V, KM F, PJ H, et al. Prevalence of Intestinal Parasites in a Low-Income Texas Community. The American journal of tropical medicine and hygiene. 2020;102(6).

3. Hotez PJ. Neglected Parasitic Infections and Poverty in the United States. 2022.

4. Liu L, Johnson HL, Cousens S, Perin J, Scott S, Lawn JE, et al. Global, regional, and national causes of child mortality: an updated systematic analysis for 2010 with time trends since 2000. Lancet. 2012;379(9832):2151–61.

5. Kotloff KL, Platts-Mills JA, Nasrin D, Roose A, Blackwelder WC, Levine MM. Global burden of diarrheal diseases among children in developing countries: Incidence, etiology, and insights from new molecular diagnostic techniques. Vaccine. 2017;35(49 Pt A):6783–9.

6. Collaborators GDD. Estimates of the global, regional, and national morbidity, mortality, and aetiologies of diarrhoea in 195 countries: a systematic analysis for the Global Burden of Disease Study 2016. Lancet Infect Dis. 2018;18(11):1211–28.

7. Shirley DT, Watanabe K, Moonah S. Significance of amebiasis: 10 reasons why neglecting amebiasis might come back to bite us in the gut. PLoS Negl Trop Dis. 2019;13(11):e0007744.

8. Turkeltaub JA, McCarty TR, Hotez PJ. The intestinal protozoa: emerging impact on global health and development. Curr Opin Gastroenterol. 2015;31(1):38–44.

9. Skappak C, Akierman S, Belga S, Novak K, Chadee K, Urbanski SJ, et al. Invasive amoebiasis: a review of Entamoeba infections highlighted with case reports. Can J Gastroenterol Hepatol. 2014;28(7):355–9.

10. Lichtenstein A, Kondo AT, Visvesvara GS, Fernandez A, Paiva EF, Mauad T, et al. Pulmonary amoebiasis presenting as superior vena cava syndrome. Thorax. 2005;60(4):350–2.

11. Wuerz T, Kane JB, Boggild AK, Krajden S, Keystone JS, Fuksa M, et al. A review of amoebic liver abscess for clinicians in a nonendemic setting. Can J Gastroenterol. 2012;26(10):729–33.

12. Petri WA, Haque R. Entamoeba histolytica brain abscess. Handb Clin Neurol. 2013;114:147–52.

13. Stanley SL. Amoebiasis. Lancet. 2003;361(9362):1025–34.

14. Haque R, Huston CD, Hughes M, Houpt E, Petri WA. Amebiasis. N Engl J Med. 2003;348(16):1565–73.

15. Houpt ER, Glembocki DJ, Obrig TG, Moskaluk CA, Lockhart LA, Wright RL, et al. The mouse model of amebic colitis reveals mouse strain susceptibility to infection and exacerbation of disease by CD4+ T cells. J Immunol. 2002;169(8):4496–503.

16. Debnath A, Rodriguez MA, Ankri S. Editorial: Recent Progresses in Amebiasis. Front Cell Infect Microbiol. 2019;9:247.

17. Mendoza Cavazos C, Knoll LJ. Entamoeba histolytica: Five facts about modeling a complex human disease in rodents. PLoS Pathog. 2020;16(11):e1008950.

18. Jyothi R, Foerster B, Hamelmann C, Shetty NP. Improved method for the concentration and purification of faecal cysts of Entamoeba histolytica for use as antigen. J Trop Med Hyg. 1993;96(4):249–50.

19. Spadafora LJ, Kearney MR, Siddique A, Ali IK, Gilchrist CA, Arju T, et al. Species-Specific Immunodetection of an Entamoeba histolytica Cyst Wall Protein. PLoS Negl Trop Dis. 2016;10(5):e0004697.

20. Ehrenkaufer GM, Suresh S, Solow-Cordero D, Singh U. High-Throughput Screening of Entamoeba Identifies Compounds Which Target Both Life Cycle Stages and Which Are Effective Against Metronidazole Resistant Parasites. Front Cell Infect Microbiol. 2018;8:276.

21. Lee KC, Lu CC, Hu WH, Lin SE, Chen HH. Colonoscopic diagnosis of amebiasis: a case series and systematic review. Int J Colorectal Dis. 2015;30(1):31–41.

22. Kantor M, Abrantes A, Estevez A, Schiller A, Torrent J, Gascon J, et al. Entamoeba Histolytica: Updates in Clinical Manifestation, Pathogenesis, and Vaccine Development. Can J Gastroenterol Hepatol. 2018;2018:4601420.

23. Mitra BN, Pradel G, Frevert U, Eichinger D. Compounds of the upper gastrointestinal tract induce rapid and efficient excystation of Entamoeba invadens. Int J Parasitol. 2010;40(6):751–60.

24. Hamano S, Asgharpour A, Stroup SE, Wynn TA, Leiter EH, Houpt E. Resistance of C57BL/6 mice to amoebiasis is mediated by nonhemopoietic cells but requires hemopoietic IL-10 production. J Immunol. 2006;177(2):1208–13.

25. Chudnovskiy A, Mortha A, Kana V, Kennard A, Ramirez JD, Rahman A, et al. Host-Protozoan Interactions Protect from Mucosal Infections through Activation of the Inflammasome. Cell. 2016;167(2):444–56.e14.

26. Ghosh SK, Van Dellen KL, Chatterjee A, Dey T, Haque R, Robbins PW, et al. The Jacob2 lectin of the Entamoeba histolytica cyst wall binds chitin and is polymorphic. PLoS Negl Trop Dis. 2010;4(7):e750.

27. Pittman KJ, Cervantes PW, Knoll LJ. Z-DNA Binding Protein Mediates Host Control of Toxoplasma gondii Infection. Infect Immun. 2016;84(10):3063–70.

28. Cervantes PW, Di Genova BM, Erazo Flores BJ, Knoll LJ. RIPK3 facilitates host resistance to oral *Toxoplasma gondii* infection. Infect Immun. 2021.

29. Sunagar R, Kumar S, Namjoshi P, Rosa SJ, Hazlett KRO, Gosselin EJ. Evaluation of an outbred mouse model for Francisella tularensis vaccine development and testing. PLoS One. 2018;13(12):e0207587.

30. Watanabe H, Numata K, Ito T, Takagi K, Matsukawa A. Innate immune response in Th1- and Th2-dominant mouse strains. Shock. 2004;22(5):460–6.

31. Ferreira BL, Ferreira É, de Brito MV, Salu BR, Oliva MLV, Mortara RA, et al. BALB/c and C57BL/6 Mice Cytokine Responses to. Front Microbiol. 2018;9:553.

32. Hartmann W, Blankenhaus B, Brunn ML, Meiners J, Breloer M. Elucidating different pattern of immunoregulation in BALB/c and C57BL/6 mice and their F1 progeny. Sci Rep. 2021;11(1):1536.

33. Denic S, Nicholls MG. Genetic benefits of consanguinity through selection of genotypes protective against malaria. Hum Biol. 2007;79(2):145–58.

34. Benton CH, Delahay RJ, Smith FAP, Robertson A, McDonald RA, Young AJ, et al. Inbreeding intensifies sex- and age-dependent disease in a wild mammal. J Anim Ecol. 2018;87(6):1500–11.

35. A A, IM A, H A, JR R, MK I. Impact of environmental conditions on the survival of cryptosporidium and giardia on environmental surfaces. Interdisciplinary perspectives on infectious diseases. 2014;2014.

36. Pawestri AR, Thima K, Leetachewa S, Maneekan P, Deesitthivech O, Pinna C, et al. Seasonal prevalence, risk factors, and One Health intervention for prevention of intestinal parasitic infection in underprivileged communities on the Thai-Myanmar border. Int J Infect Dis. 2021;105:152–60.

37. Jaran AS. Prevalence and seasonal variation of human intestinal parasites in patients attending hospital with abdominal symptoms in northern Jordan. East Mediterr Health J. 2017;22(10):756–60.

38. Chowdhury FR, Ibrahim QSU, Bari MS, Alam MMJ, Dunachie SJ, Rodriguez-Morales AJ, et al. Correction: The association between temperature, rainfall and humidity with common climate-sensitive infectious diseases in Bangladesh. PLoS One. 2020;15(4):e0232285.

39. Turnbaugh PJ, Hamady M, Yatsunenko T, Cantarel BL, Duncan A, Ley RE, et al. A core gut microbiome in obese and lean twins. Nature. 2009;457(7228):480–4.

40. McNulty NP, Yatsunenko T, Hsiao A, Faith JJ, Muegge BD, Goodman AL, et al. The impact of a consortium of fermented milk strains on the gut microbiome of gnotobiotic mice and monozygotic twins. Sci Transl Med. 2011;3(106):106ra.

41. Walderich B, Müller L, Bracha R, Knobloch J, Burchard GD. A new method for isolation and differentiation of native Entamoeba histolytica and E. dispar cysts from fecal samples. Parasitol Res. 1997;83(7):719–21.

42. Morgulis A, Coulouris G, Raytselis Y, Madden TL, Agarwala R, Schäffer AA. Database indexing for production MegaBLAST searches. Bioinformatics. 2008;24(16):1757–64.

43. Guindon S, Dufayard JF, Lefort V, Anisimova M, Hordijk W, Gascuel O. New algorithms and methods to estimate maximum-likelihood phylogenies: assessing the performance of PhyML 3.0. Syst Biol. 2010;59(3):307–21.

